# Metabolic reprogramming of immune cells in HIV infection and treatment

**DOI:** 10.1101/2024.09.26.615098

**Authors:** Phoebe Xu, Di Wu

**Affiliations:** Enloe Magnet High School, Raleigh, North Carolina 27610, USA; Department of Biostatistics, Gillings School of Global Public Health, University of North Carolina at Chapel Hill, Chapel Hill, NC 27599, USA

**Keywords:** Single cell, scRNA-seq, metabolite pathway, HIV, ART, metabolites-mediated cell-cell communication

## Abstract

**BACKGROUND:** Immune cells undergo metabolic reprogramming, altering their energy production in response to HIV infection. During acute HIV infection, CD4+ T cells increase glycolysis to fuel viral replication and defense, but prolonged activation leads to immune exhaustion, with reduced T cell function and impaired pathogen elimination, contributing to AIDS progression. However, the details Metabolic dynamics of Immune Cells during HIV Infection (UT) and after receiving antiretroviral therapy (ART) remains unknown. Communication between Immune cells will be modified to response HIV infection, which can be mediated by metabolites. In this study, we deciphered Metabolic pathway dynamics of all Immune Cells by comparing UT and ART with seronegative controls (SN). Additionally, we investigate alteration of metabolite-mediated cell-cell communications in HIV Infection.

**METHODS:** Single cell RNA sequencing (scRNA-seq) data of PBMC from SN, UT and ART were analyzed with KEGG Metabolic pathways by irGSEA and algorithm MEBOCOST to study cell-cell communications mediated by metabolites from sender cells and sensor proteins on receiver cells.

**RESULTS:** lots of differentially expressed Metabolic Pathways were identified for immune cells, including Energy Production and Utilization, Lipid Metabolism, Amino Acid Metabolism, Glycosylation and Glycan Biosynthesis and Oxidative Stress and Antioxidant Defense.

**CONCLUSIONS:** The analysis highlights the intricate metabolic interplay between monocytes and other immune cells, reflecting their central role in orchestrating immune responses through metabolite exchange. These insights may open avenues for therapeutic interventions, targeting metabolic pathways to enhance immune function or inhibit pathogen survival.

## 1. Introduction

Immune cells undergo metabolic reprogramming, altering their energy production in response to infection^1–3^. This is especially significant in chronic infections like HIV, affecting disease progression and immune function^4^. During infection, activated immune cells, particularly T cells and macrophages, shift from oxidative phosphorylation (OXPHOS) to glycolysis to meet higher energy demands. This metabolic shift, known as the “Warburg effect” in T cells, supports rapid proliferation, cytokine production, and pathogen response^5^.

HIV infection drives chronic immune activation, forcing immune cells, especially CD4+ T cells, to continually alter their metabolism^4,6,7^. During acute HIV infection, CD4+ T cells increase glycolysis to fuel viral replication and defense, but prolonged activation leads to immune exhaustion, with reduced T cell function and impaired pathogen elimination, contributing to AIDS progression^8^. Similarly, macrophages shift toward glycolysis when exposed to inflammation, enhancing their role as viral reservoirs and complicating immune clearance of HIV.

Single-cell RNA sequencing (scRNA-seq) has revolutionized our understanding of cellular heterogeneity and the intricate changes that occur at the transcriptomic level in various diseases, especially SARS-CoV-2 ^9–13^. Some case studies of scRNA-seq in HIV infection was reported^14–16^. This technology allows for the dissection of complex cellular populations and the identification of distinct metabolic states within individual cells. In the context of HIV, scRNA-seq provides critical insights into how the virus alters the metabolic landscape of immune cells, contributing to immune dysfunction and disease progression. This comprehensive approach allowed researchers to compare the metabolic changes across different stages of HIV infection and treatment. Unfortunately, there have few of studies focus on decipher dynamics of metabolic states for immune cells during HIV infection and treatment.

In this study, we will use scRNA-seq to analyze peripheral blood immune cells from three groups: chronic untreated HIV-1 individuals (UT), HIV-1-infected individuals receiving antiretroviral therapy (ART), and seronegative controls (SN)). We grouped cells into sub cell types. Then we comprehensively compare all metabolic pathways among SN, UT and ART for each sub cell types. Moreover, a new computational algorithm called MEBOCOST (Metabolite-mediated Cell Communication Modeling by Single Cell Transcriptome) are used to quantitatively infer metabolite-mediated intercellular communications^17^. Lots of cell-cell communications with big change of metabolites and sensor are identified. Insights into these metabolic shifts open avenues for therapeutic interventions, targeting metabolic pathways to enhance immune function or inhibit pathogen survival.

## 2. Materials and methods

Description of patients and donors. Deidentified peripheral blood mononuclear cell (PBMC) samples were used from existing cohorts of HIV-1-infected and seronegative individuals stored at the Duke Human Vaccine Institute that were approved by the Duke Medicine Institutional Review Boards as well as the ethics boards of the local sites^5^. Cells from six untreated HIV-infected, three ART HIV-1-infected (HIV-1 viral load below the limit of detection), and three HIV-1 seronegative (control) individuals were used to perform single-cell RNA sequencing studies (10× Genomics;).

### scRNA-seq data analysis

Raw fastq were download from NCBI SRA database. Then, CellRanger software (version 6.01, 10X Genomics) was used to preprocess fastq files and count gene expression matrices for each sample. Gene expression matrices were input to R package Seurat (version 4.3.0) for downstream analyses^18^. Briefly, scRNA-seq data from different samples were done quality control (QC), filtered out low expression genes and low-quality cells, then merged. The distribution of UMI, all genes, mitochondrial genes, and ribosomal genes of each sample was calculated. Low-quality cells with <200 UMIs or with a proportion of mitochondrial genes >5% were removed. The first 30 principal components, along with the first 3,000 highly variable genes, were acquired for further analysis. The disturbance of UMI counts as well as the proportion of mitochondrial UMI counts were regressed with “*ScaleData*” function. Cell cycle score was calculated, and cell cycle effect was removed as well. Thereafter, the major cell types were clustered utilizing “*FindClusters*” function, which was visualized with uniform manifold approximation and projection (UMAP) to reduce data dimensions. Through marker genes of each cell, a cluster was calculated using “*FindAllMarkers*” function. The main cell types were identified based on predicted and known marker genes acquired from the SingleR (https://github.com/LTLA/SingleR<u>) and</u> CellMarker database (http://biocc.hrbmu.edu.cn/CellMarker/) ^19^.

### Sub-cell type analysis

Raw gene expression count of cells for sub-cell type were extracted from previous cleaned data and merged to a Seurat object using the Seurat. Gene expression matrices were normalized to total cellular read count, and highly variably genes (HVG) selected from the normalized data using Seurat SCTransform function with default parameters. Batch effects among different samples were eliminated utilizing “Harmony”^20^ because batch effects were observed after all samples data were integrated. Following dimensional reduction and clustering used same procedure for whole cell population.

### Metabolic pathway analysis

Metabolic pathway was download from KEGG database^9^. Metabolic pathway for each sub cell type between SN, UT and ART are analyzed using tool-“irGSEA” ^21^. First, all cells from each cell type (e.g. T and NK cell are regrouped into “sub cell type + Treatment”, e.g. “Tem_SN”. Comparison of genes expression change between different combination of sub cell type and treatment were calculated via “FindMarkers” function. “ssGSEA” was carried out to detect enriched metabolic pathway ^22^. The terms with a p-value <0.05 were regarded as significant enrichment.

### Metabolite-based Cell-cell communication analysis

Algorithm MEBOCOST was used to study cell-cell communications mediated by metabolites from sender cells and sensor proteins on receiver cells. The data was analyzed following the tutorial on the MEBOCOST website (https://github.com/ kaifuchenlab/MEBOCOST). Note that we don’t conduct metabolomics data analysis in this study. Instead, scRNA-seq expression data was used to estimate metabolite abundance and calculate communication score for each condition.

### Statistical analysis

Statistical analysis was implemented utilizing R software (version 4.3.2; https://www.r-project.org/). Comparisons between groups were analyzed with students’ t-test or one-way analysis of variance. P<0.05 indicated statistical significance.

### Data availability

scRNA-Seq data in this study were downloaded from the NCBI SRA database under the BioProject ID: PRJNA681021.

## 3. Results

### 3.1 scRNA-seq identifies diverse cell types of PBMC in HIV infection and treatment

For an unbiased analysis of all metabolic pathways reprogrammed by HIV, scRNA-seq analysis of PBMC isolated from 3 seronegative (SN), 6 untreated (UT) and 3 antiretroviral therapy (ART) human patients were performed. We identified a diverse range of cell types, including T cells, B cell, Monocytes cells, and other cells (Figure 1A). Known cell markers facilitated annotation of cell types, such as CD3D and CD3E for T cells, MS4A1, CD79A and CD79B for B cells, Cd14 and FCGR3A for monocytes (Figure 1B). Thereafter, we displayed the distribution of the above cell types in PBMC from SN, UT and ART (Figure 1C), respectively.

**Figure 1.**
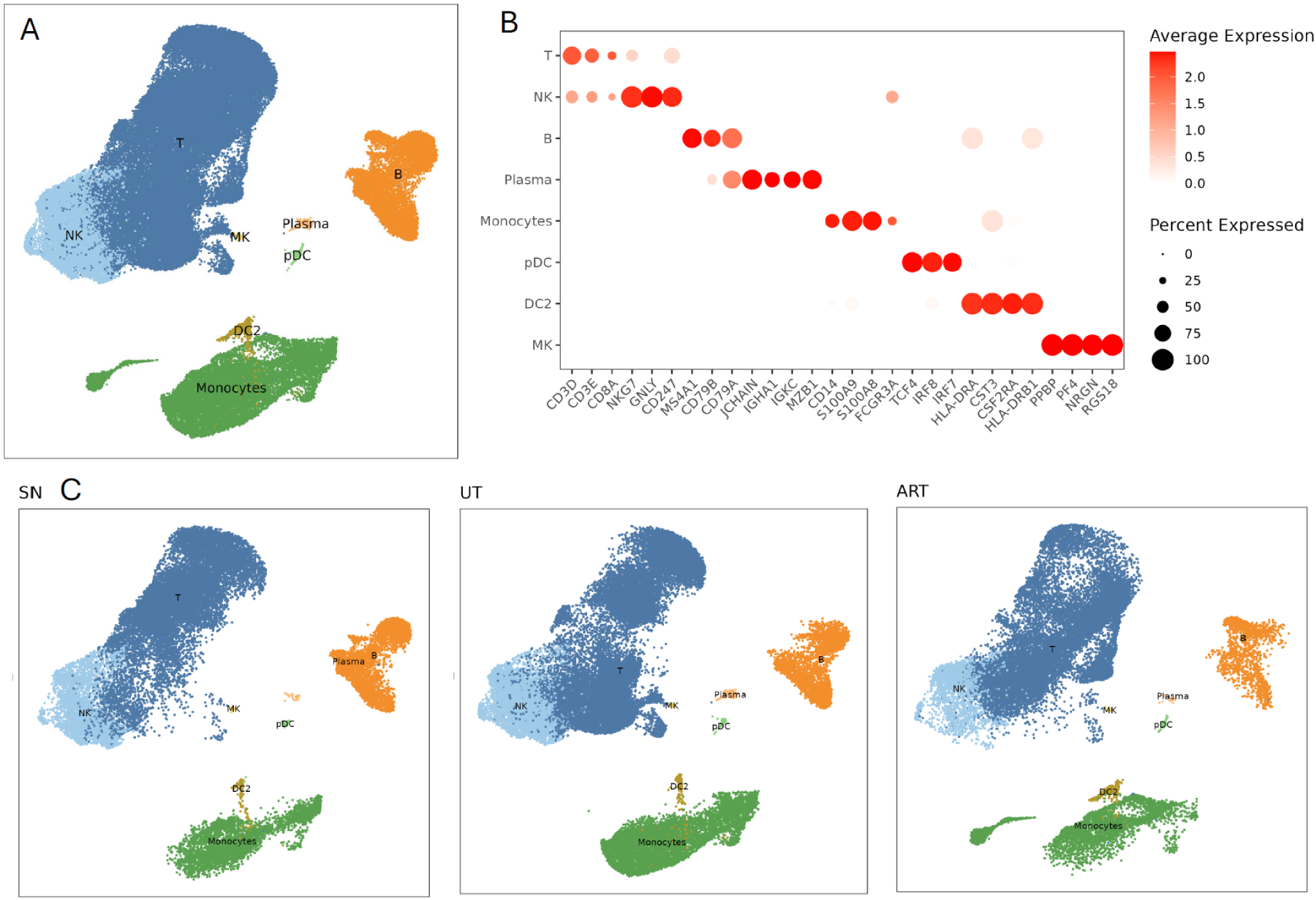
scRNA-seq analysis of human PBMC from 3 SN (seronegative), 6 UT (untreated) patients and 3 ART (antiretroviral therapy), identifies distinct cell types. (A) UMAP plot of 8 cell types of PBMC. (B) Dotplot showing the gene expression of marker genes of each cell type. The size of dots indicates the abundance level, and color indicates the expression level. (C) UMAP plots split by SN, UT and ART. NK: natural killer cell, MK: megakaryocyte, pDC: plasma dendritic cell.

### 3.2 Single-cell RNA-seq revealed that T cell reprogramed their metabolic pathway to response HIV infection

It was observed that under normal, non-infectious conditions, immune cells such as macrophages and dendritic cells utilize OXPHOS as their primary energy source. However, in response to infection, many immune cells, particularly activated T cells and macrophages, shift to glycolysis to meet the increased energy demands of mounting an immune response. To investigate the metabolic pathway dynamics of T cells in response to HIV infection, we isolated T and NK cell populations, further subdivided them into distinct subtypes, and examined the heterogeneity across these subpopulations (Figure 2A). Multiple well-known T subpopulations were recaptured, including CD4+ T, CD8+ T and NK. We combined markers reported in the literature and predicted using “Findmarker” to annotate the subtypes (Figure S1B). The composition of these sub-T and NK populations showed differences between SN, UT and ART patients (Figure 2B, Figure S1A). Compared to SN, the cell proportions of CD8_Treg, Tem, CD8_Temra1, CD8_Temra2 and CD8_Temra3 were increased in UT patients, while the cell proportions of CD4-Treg, CD4-Tcm, and Mast were decreased in UT patients. Additionally, CD8-Tcm, CD8_Temra2, and CD8_Temra3 were increased in ART samples.

**Figure 2.**
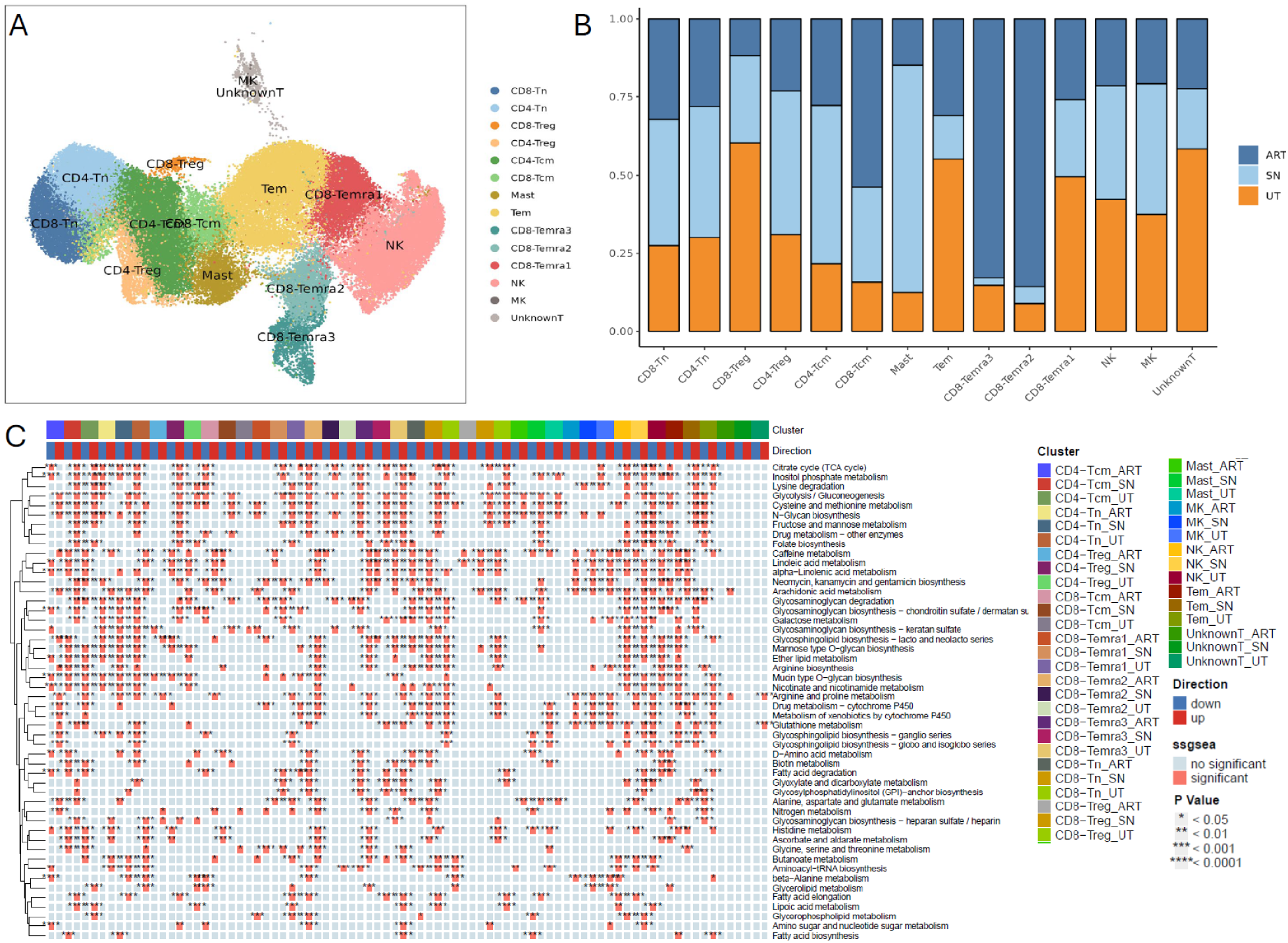
Identification of subpopulations of T and NK cells of PBMC and dynamic of their metabolism pathways between SN, UT and ART. A. UMAP plot of 14 sub-T and NK cell types of PBMC. B. The fraction of each sub-T cell type that originated from SN, UT, ART samples. C. Metabolism Pathways in T Cell Subtypes from HIV and ART Patients. This heatmap visualizes the top 50 metabolism-related pathways identified by irGSEA (immune-related Gene Set Enrichment Analysis) across various T cell subtypes in SN, UT and ART patients. The analyzed subtypes include CD4+ and CD8+ T cells in various states such as Tcm, Tn, Treg, and terminally differentiated effector memory (Temra). These cells were further grouped by treatment conditions: ART, SN, and UT. The heatmap presents the differential expression patterns for each T cell subtype under different treatment conditions. Pathways are upregulated (marked in red) or downregulated (marked in blue) based on significance levels, with * indicating different p-value thresholds (e.g., *p* < 0.05, \**p* < 0.01, etc.). Tn: naïve T cell, Tcm: central memory T cell, Treg: regulatory T cell, Tem: effector memory T cells, Temra: terminally differentiated effector memory T cells.

We then conducted metabolism-related pathways analysis using irGSEA (The integration of single cell rank-based gene set enrichment analysis). Top 50 metabolism-related pathways across various T cell subtypes in SN, UT and ART patients identified by irGSEA were visualized with heatmap (Figure 2C). The top enriched pathways include those involved in the Citrate cycle (TCA), glycolysis/gluconeogenesis, lipid metabolism, and amino acid metabolism (e.g., glycine, serine, and threonine metabolism). These pathways show varying levels of activity across T cell subtypes. Metabolism in T cells is significantly altered in response to HIV infection and ART.

The metabolic reprogramming of T and NK cells during HIV infection reveals significant alterations in various metabolic pathways, which can impact immune cell function and overall immune response. Here are the key findings and their potential implications. For CD4-Treg, D-Amino acid metabolism, beta-Alanine metabolism, Glycerolipid metabolism, and Lipoic acid metabolism were up-regulated on UT patients. These changes suggest enhanced amino acid utilization, lipid synthesis, and antioxidant defense. While Ether lipid metabolism, Nicotinate and nicotinamide metabolism, Glutathione metabolism, and Biotin metabolism were down-regulated on UT patients. This indicates reduced lipid signaling, NAD+ synthesis, antioxidant capacity, and biotin-dependent carboxylation reactions.

For CD8-Treg, Fatty acid degradation, Glyoxylate and dicarboxylate metabolism, Fatty acid elongation, and Lipoic acid metabolism were up-regulated on UT patients. These pathways support increased energy production, lipid synthesis, and antioxidant defense. While, Glycosaminoglycan degradation, Glycosphingolipid biosynthesis (lacto and neolacto series), Ether lipid metabolism, Arginine biosynthesis, Arginine and proline metabolism, Drug metabolism (cytochrome P450), Histidine metabolism, Ascorbate and aldarate metabolism were down-regulated on UT patients. This suggests reduced structural integrity, signaling, amino acid synthesis, and detoxification processes.

For Tem, Histidine metabolism, Fatty acid degradation, D-Amino acid metabolism, Metabolism of xenobiotics by cytochrome P450, Mucin type O-glycan biosynthesis, Nicotinate and nicotinamide metabolism, Glycosphingolipid biosynthesis (lacto and neolacto series) were up-regulated on UT patients. These changes indicate enhanced amino acid utilization, lipid degradation, detoxification, and glycan synthesis. While, Glycolysis/Gluconeogenesis, Cysteine and methionine metabolism, N-Glycan biosynthesis, Fructose and mannose metabolism, Drug metabolism (other enzymes), Folate biosynthesis, Glycosaminoglycan degradation, Arginine biosynthesis, Glycosylphosphatidylinositol (GPI)-anchor biosynthesis, Nitrogen metabolism, Glycosaminoglycan biosynthesis (heparan sulfate/heparin) were down-regulated on UT patients. This suggests reduced energy production, amino acid synthesis, glycan synthesis, and structural integrity.

For CD8-Temra1, Mannose type O-glycan biosynthesis, Metabolism of xenobiotics by cytochrome P450 were up-regulated on UT patients. These changes support enhanced glycan synthesis and detoxification. While, Lysine degradation, Glycolysis/Gluconeogenesis, Cysteine and methionine metabolism, N-Glycan biosynthesis, Fructose and mannose metabolism, Folate biosynthesis, Arachidonic acid metabolism, Glycosaminoglycan biosynthesis (keratan sulfate), Arginine and proline metabolism, Glycosphingolipid biosynthesis (ganglio and globo/isoglobo series), Glycerolipid metabolism, Fatty acid elongation, Lipoic acid metabolism, Glycerophospholipid metabolism were down-regulated on UT patients. This indicates reduced amino acid degradation, energy production, lipid synthesis, and antioxidant defense.

For CD8-Temra2, Nicotinate and nicotinamide metabolism were up-regulated on UT patients. This suggests increased NAD+ synthesis. While, Citrate cycle (TCA cycle), Inositol phosphate metabolism, Lysine degradation, Glycolysis/Gluconeogenesis, Cysteine and methionine metabolism, N-Glycan biosynthesis, Fructose and mannose metabolism, Drug metabolism (other enzymes) were down-regulated on UT patients. This indicates reduced energy production, signaling, amino acid degradation, and detoxification.

For CD8-Temra3, Glycine, serine and threonine metabolism, Nicotinate and nicotinamide metabolism, Arginine and proline metabolism were up-regulated on UT patients. These changes support enhanced amino acid synthesis and NAD+ production. While, Mannose type O-glycan biosynthesis, Ether lipid metabolism, Mucin type O-glycan biosynthesis, Biotin metabolism, Nitrogen metabolism, Glycosaminoglycan biosynthesis (heparan sulfate/heparin) were down-regulated on UT patients. This suggests reduced glycan synthesis, lipid signaling, biotin-dependent reactions, and structural integrity.

The observed metabolic reprogramming in T and NK cells during HIV infection suggests several functional implications. For CD4-Treg, D−Amino acid metabolism, beta−Alanine metabolism, Glycerolipid metabolism and Lipoic acid metabolism were upregulated on UT patients, while Ether lipid metabolism, Nicotinate and nicotinamide metabolism, Glutathione metabolism and Biotin metabolism were down-regulated.

For CD8-Treg, Fatty acid degradation, Glyoxylate and dicarboxylate metabolism, Fatty acid elongation and Lipoic acid metabolism were upregulated on UT patients, while Glycosaminoglycan degradation, Glycosphingolipid biosynthesis − lacto and neolacto series, Ether lipid metabolism, Arginine biosynthesis, Arginine and proline metabolism, Drug metabolism − cytochrome P450, Metabolism of xenobiotics by cytochrome P450, Glycosaminoglycan biosynthesis − heparan sulfate / heparin, Histidine metabolism, Ascorbate and aldarate metabolism were down-regulated.

For Tem, Histidine metabolism, Fatty acid degradation, D−Amino acid metabolism, Metabolism of xenobiotics by cytochrome P450, Mucin type O−glycan biosynthesis, Nicotinate and nicotinamide metabolism, Glycosphingolipid biosynthesis − lacto and neolacto series were upregulated on UT patients, while Glycolysis / Gluconeogenesis, Cysteine and methionine metabolism, N−Glycan biosynthesis, Fructose and mannose metabolism, Drug metabolism − other enzymes, Folate biosynthesis, Glycosaminoglycan degradation, Arginine biosynthesis, Glycosylphosphatidylinositol (GPI)−anchor biosynthesis, Nitrogen metabolism and Glycosaminoglycan biosynthesis – heparan sulfate / heparin were down-regulated.

For CD8-Temra1, Mannose type O−glycan biosynthesis, Metabolism of xenobiotics by cytochrome P450 were upregulated on UT patients, while Lysine degradation, Glycolysis / Gluconeogenesis, Cysteine and methionine metabolism, N−Glycan biosynthesis, Fructose and mannose metabolism, Folate biosynthesis, Arachidonic acid metabolism, Glycosaminoglycan biosynthesis − keratan sulfate, Arginine and proline metabolism, Glycosphingolipid biosynthesis − ganglio series, Glycosphingolipid biosynthesis − globo and isoglobo series, Glycerolipid metabolism, Fatty acid elongation, Lipoic acid metabolism and Glycerophospholipid metabolism were down-regulated.

For CD8-Temra2, Nicotinate and nicotinamide metabolism were upregulated on UT patients, while Citrate cycle (TCA cycle), Inositol phosphate metabolism, Lysine degradation, Glycolysis / Gluconeogenesis, Cysteine and methionine metabolism, N−Glycan biosynthesis, Fructose and mannose metabolism and Drug metabolism − other enzymes were down-regulated.

For CD8-Temra2, Glycine, serine and threonine metabolism, Nicotinate and nicotinamide metabolism, Arginine and proline metabolism were upregulated on UT patients, while Mannose type O−glycan biosynthesis, Ether lipid metabolism, Mucin type O−glycan biosynthesis, Biotin metabolism, Nitrogen metabolism and Glycosaminoglycan biosynthesis − heparan sulfate / heparin were down-regulated.

ART groups show unique enrichment patterns in metabolic pathways compared to untreated and seronegative individuals. Pathways like glycolysis, lipid biosynthesis, and amino acid degradation are particularly modulated in CD4+ and CD8+ T cells, suggesting shifts in energy and nutrient utilization. In ART (CD4-Tcm_ART) cells, glycolysis/gluconeogenesis shows a marked upregulation, suggesting that these cells rely on glycolytic pathways for energy production while on therapy. This could reflect an enhanced metabolic state to support cell survival and function under ART. In contrast, untreated (CD4-Tcm_UT) and seronegative (CD4-Tcm_SN) groups exhibit downregulation of this pathway, implying that the metabolic demand in these cells may be lower in the absence of ART or viral infection. CD8-Temra1_ART cells show significant upregulation of the TCA cycle, indicating increased mitochondrial oxidative phosphorylation activity in ART cells. This suggests enhanced ATP production through aerobic respiration, possibly contributing to long-term survival and effector functions. The CD8-Temra1_SN and CD8-Temra1_UT groups, on the other hand, show reduced TCA cycle activity, potentially reflecting a lower metabolic demand or a shift towards glycolysis in the absence of ART. Arginine and proline metabolism is significantly upregulated in CD4-Treg_ART cells. Arginine metabolism is crucial for Treg function and immune regulation, suggesting that ART enhances the metabolic processes that sustain Treg-mediated immune suppression. CD4-Treg_UT and CD4-Treg_SN groups display less significant changes in this pathway, potentially indicating that regulatory T cells in untreated or seronegative patients are metabolically less active or regulated differently. In CD8-Tn_ART cells, glutathione metabolism is highly upregulated, indicating enhanced antioxidant defense mechanisms. Glutathione plays a key role in protecting cells from oxidative stress, which may be elevated in ART patients due to the immune activation and metabolic stress. The pathway is less active in CD8-Tn_SN and CD8-Tn_UT, suggesting that oxidative stress responses are less pronounced in these populations, possibly due to lower immune activation. In NK_ART cells, fatty acid biosynthesis is significantly upregulated, implying that these cells may shift towards lipid synthesis under ART to support membrane biosynthesis and cellular growth. Conversely, the NK_SN and NK_UT groups show a relative downregulation of fatty acid biosynthesis, indicating a lower demand for lipid production, possibly due to reduced activation or proliferative needs. Mast cells in the ART group (Mast_ART) show significant upregulation of glycosphingolipid biosynthesis, particularly in the ganglio series. Glycosphingolipids are important for cell membrane stability and signal transduction, suggesting that ART influences lipid-based signaling pathways in mast cells. This pathway is notably less active in Mast_SN and Mast_UT, which may suggest that HIV infection or ART modulates glycosphingolipid biosynthesis specifically in mast cells.

### 3.3 Monocytes and DC reprogramed their metabolic pathway to response HIV infection

To investigate the metabolic pathway dynamics of monocytes and dendritic cells (DCs) in response to HIV infection, we extracted these populations and further subdivided them into distinct subclusters to assess heterogeneity among subpopulations (Figure 3A). Several well-known monocyte and DC subpopulations were identified, including classical monocytes (e.g., C-Mono1), non-classical monocytes (e.g., NC-Mono), and DCs (e.g., pDC). Subtypes were annotated using a combination of literature-reported markers and those predicted by the "Findmarker" tool (Figure S2B). The composition of these monocyte subpopulations varied between SN, UT, and ART patients (Figure 3B, Figure S2A). Compared to SN, the cell proportion of C1QA-Mono was increased in UT samples. Additionally, the cell proportions of C1QA-Mono, pDC, CD2, and IFITM3-Mono were increased in ART samples.

**Figure 3.**
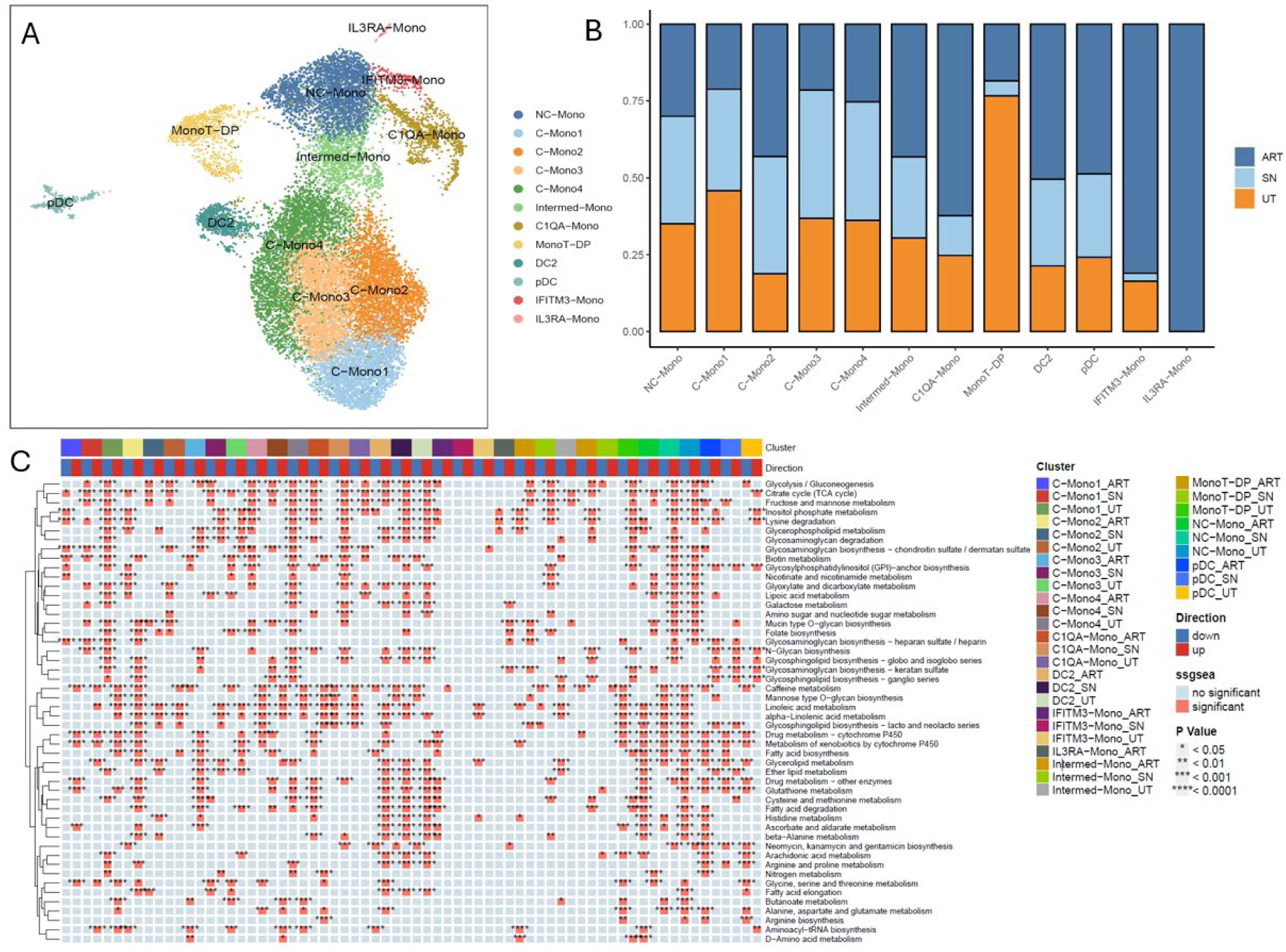
Identification of subpopulations of Monocytes and DC cells of PBMC and dynamic of their Metabolism Pathways between SN, UT and ART. A. UMAP plot of 12 sub-Monocytes and DC cell types of PBMC. B. The fraction of each sub-Monocytes and DC cell type that originated from SN, UT, ART samples. C. Metabolism Pathways in sub-Monocytes and DC from SN, UT and ART Patients. This heatmap shows the top 50 metabolism-related pathways identified by irGSEA (immune-related Gene Set Enrichment Analysis) in various monocyte subtypes from HIV and ART patients. The monocyte subtypes analyzed include classical monocytes (C-Mono), intermediate monocytes (Intermed-Mono), non-classical monocytes (NC-Mono), and plasmacytoid dendritic cells (pDC), as well as specialized monocyte subsets like IFITM3-Mono and C1QA-Mono. These subtypes are grouped under different conditions: ART (antiretroviral therapy), SN (seronegative), and UT (untreated). Pathways are either upregulated (marked in red) or downregulated (marked in blue) across different monocyte subtypes and conditions. The color intensity represents the level of significance, indicated by p-values (*p* < 0.05, \**p* < 0.01, etc.).C-Mono: classical monocytes, NC-Mono: non-classical monocytes, Intermed-mono: intermediate monocytes.

Similar to the analysis performed for T cell subtypes, we conducted a metabolism-related pathway analysis for monocyte and dendritic cell (DC) subtypes using irGSEA. The pathways examined span various metabolic processes, including glycolysis/gluconeogenesis, the citric acid cycle (TCA), fatty acid metabolism, amino acid metabolism, nucleotide sugar metabolism, glutathione metabolism, and glycosylation pathways (e.g., N-glycan biosynthesis). The top 50 metabolism-related pathways across different monocyte and DC subtypes in SN, UT, and ART patients, as identified by irGSEA, are visualized in a heatmap (Figure 3C).

Here are the key findings and their potential implications. For C-Mono1, Pathways involved in lipid metabolism (e.g., Glycerolipid metabolism, Ether lipid metabolism), amino acid metabolism (e.g., Cysteine and methionine metabolism), and nucleotide metabolism (e.g., Inositol phosphate metabolism) are down-regulated on UT. This suggests a reduced capacity for membrane synthesis, signaling, and energy production. Pathways such as Mannose type O-glycan biosynthesis, Fatty acid biosynthesis, and Glutathione metabolism are up-regulated on UT, indicating an increased focus on glycan synthesis, lipid production, and antioxidant defense.

For C-Mono2, Glycosaminoglycan biosynthesis and amino sugar metabolism are down-regulated on UT, which may affect cell structure and signaling. Fatty acid biosynthesis is up-regulated on UT and ART, suggesting enhanced lipid synthesis for membrane formation and energy storage.

For Intermed-Mono, Pathways involved in lipid metabolism (e.g., Glycerolipid metabolism, Ether lipid metabolism) and glycosphingolipid biosynthesis are down-regulated on UT, indicating reduced lipid synthesis and signaling. Mucin type O-glycan biosynthesis is up-regulated on UT and ART, which may enhance cell-cell interactions and immune recognition.

For C1QA-Mono, Lipoic acid metabolism is up-regulated on UT, suggesting increased antioxidant defense. Pathways like Caffeine metabolism and Linoleic acid metabolism show Up-regulated on ART, but down-regulated on UT and SN, indicating varying metabolic demands under different conditions. Glycolysis/Gluconeogenesis and Citrate cycle (TCA cycle) are down-regulated on ART, but up-regulated on UT and SN, affecting energy production and metabolic flexibility.

For IFITM3-Mono, Glycosaminoglycan biosynthesis is up-regulated on UT, enhancing structural integrity and signaling. Pathways like Linoleic acid metabolism and Glycerolipid metabolism are down-regulated on UT, reducing lipid synthesis and signaling.

For pDC, Pathways such as Citrate cycle (TCA cycle) and Inositol phosphate metabolism are up-regulated on UT, indicating increased energy production and signaling. Down-regulated on UT: Pathways like Arachidonic acid metabolism and Glycine, serine, and threonine metabolism are down-regulated on UT, potentially reducing inflammatory responses and amino acid synthesis.

### 3.4 Single-cell RNA-seq revealed that metabolic pathways were reprogramed in B and plasma cell to response HIV infection

To study metabolic pathway dynamic of B and plasma cell to response HIV infection, we extracted B and plasma populations and then further subclustered to investigate their heterogeneity among subpopulations (Figure 4A). Multiple well-known B and plasma subpopulations were recaptured, including B-naïve (e.g. Bn-1), B-median (e.g. Bm-1) and plasma. We combined markers reported in the literature and predicted using “Findmarker” to annotate the subtypes (Figure S3B). The composition of these sub-B and plasma populations showed differences between SN, UT and ART patients (Figure 4B, Figure S3A). Compared to SN, the cell proportions of Plasma, age-B, and Bn3 were increased in UT samples, while Bn2 was increased in ART samples.

**Figure 4.**
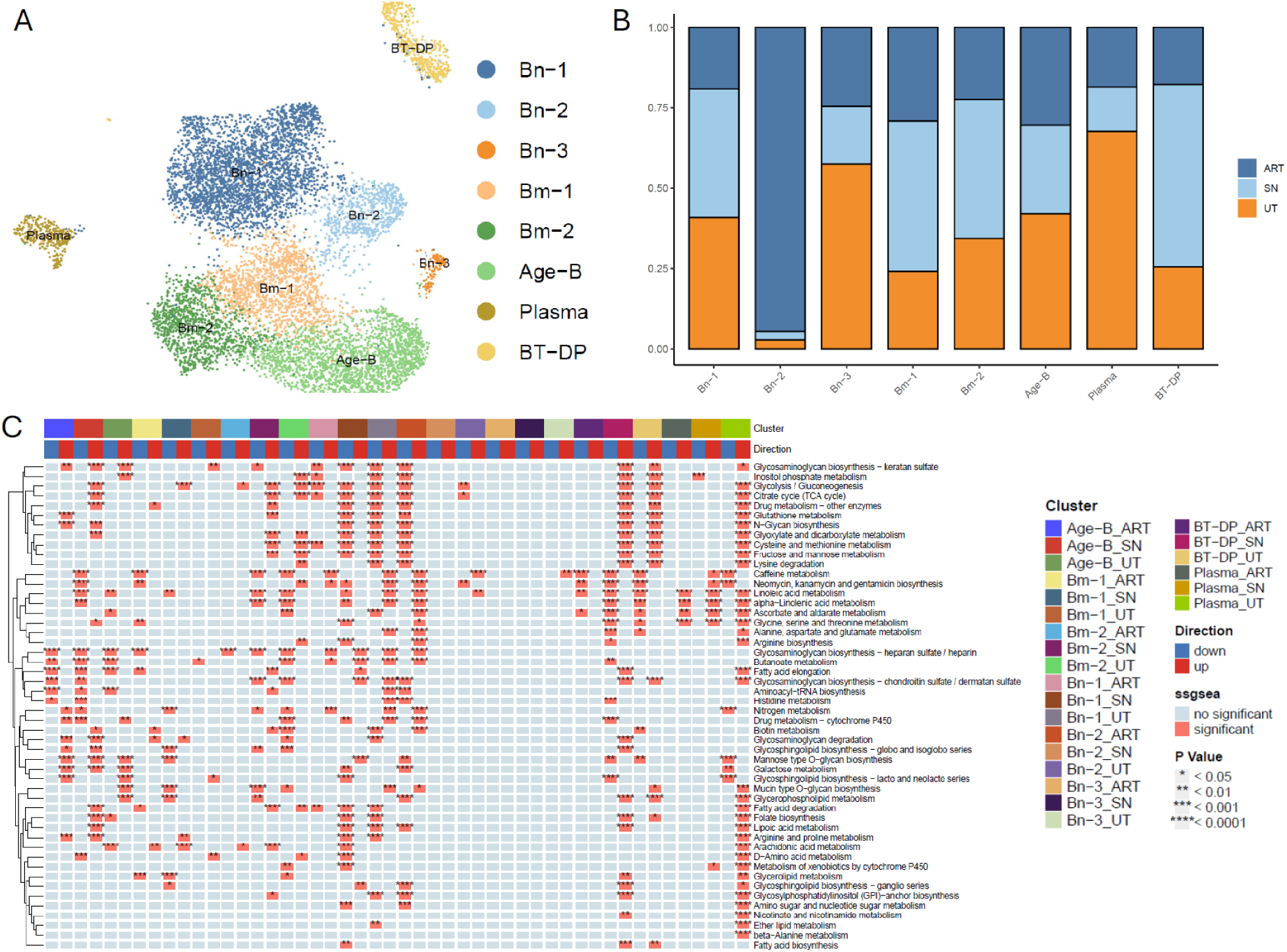
Identification of subpopulations of B and plasma cells of PBMC and dynamic of their metabolism pathways between SN, UT and ART. A. UMAP plot of 8 sub-B and plasma cell types of PBMC. B. The fraction of each sub-B and plasma cell type that originated from SN, UT, ART samples. C. Metabolism Pathways in sub-B and plasma Cell Subtypes from HIV and ART Patients. This heatmap represents the activity of various metabolic pathways in B cells from SN, UT, and ART groups. The pathways were analyzed using the irGSEA method, and the significance of regulation is indicated by asterisks (*p* < 0.05, p < 0.01, *p* < 0.001, p < 0.0001). Bn: naïve B cell; Bm: memory B cell; BT-DP: B and T double positive cell.

We then conducted metabolism-related pathways analysis using irGSEA (immune-related Gene Set Enrichment Analysis). Top 50 metabolism-related pathways across various B and plasma cell subtypes in SN, UT and ART patients identified by irGSEA were visualized with heatmap (Figure 4C). Key metabolic pathways showing significant changes include glycine, serine, and threonine metabolism, glycolysis/gluconeogenesis, citrate cycle (TCA cycle), glutathione metabolism, and several others. The direction of regulation (up or down) is also indicated, providing insights into the metabolic reprogramming of B cells in response to HIV infection and ART treatment.

The heatmap analysis of metabolic pathways in B cell scRNA-seq data from HIV patients undergoing different treatment conditions (SN, UT, ART) reveals significant changes in various metabolic processes.

The differential regulation of metabolic pathways across B cell subtypes in HIV patients under various conditions (SN, UT and ART) shows distinct patterns. In Age-B subtypes, key pathways like glycolysis/gluconeogenesis, citrate cycle (TCA cycle) and “glycine, serine and threonine metabolism” display notable differences between SN and ART/UT, with ART and UT showing a downregulation in these pathways. Bm-1 and Bm-2 subtypes exhibit differential regulation of lipid metabolism, such as fatty acid elongation and degradation, with ART leading to suppression, while UT shows more active metabolism. Bn-1 to Bn-3 subtypes, especially under UT and ART, demonstrate downregulation of amino acid metabolism pathways, including glycine, serine, and threonine metabolism. These findings suggest that ART still under impact of metabolic reprogramming differently across B cell subtypes, influencing immune function.

### 3.5 Single-cell RNA-seq revealed that regulation of immune cell metabolism and efflux pathways in HIV infection

Major sub cell types depicted include classical monocytes (C-Mono), intermediate monocytes (Intermed-Mono), non-classical monocytes (NC-Mono), CD4+ and CD8+ T cells (Tcm, Tem, Temra), B cells (Bn-1, Bn-2), NK cells, and dendritic cells (pDC, DC2) (Figure 5A, Figure S4).

**Figure 5.**
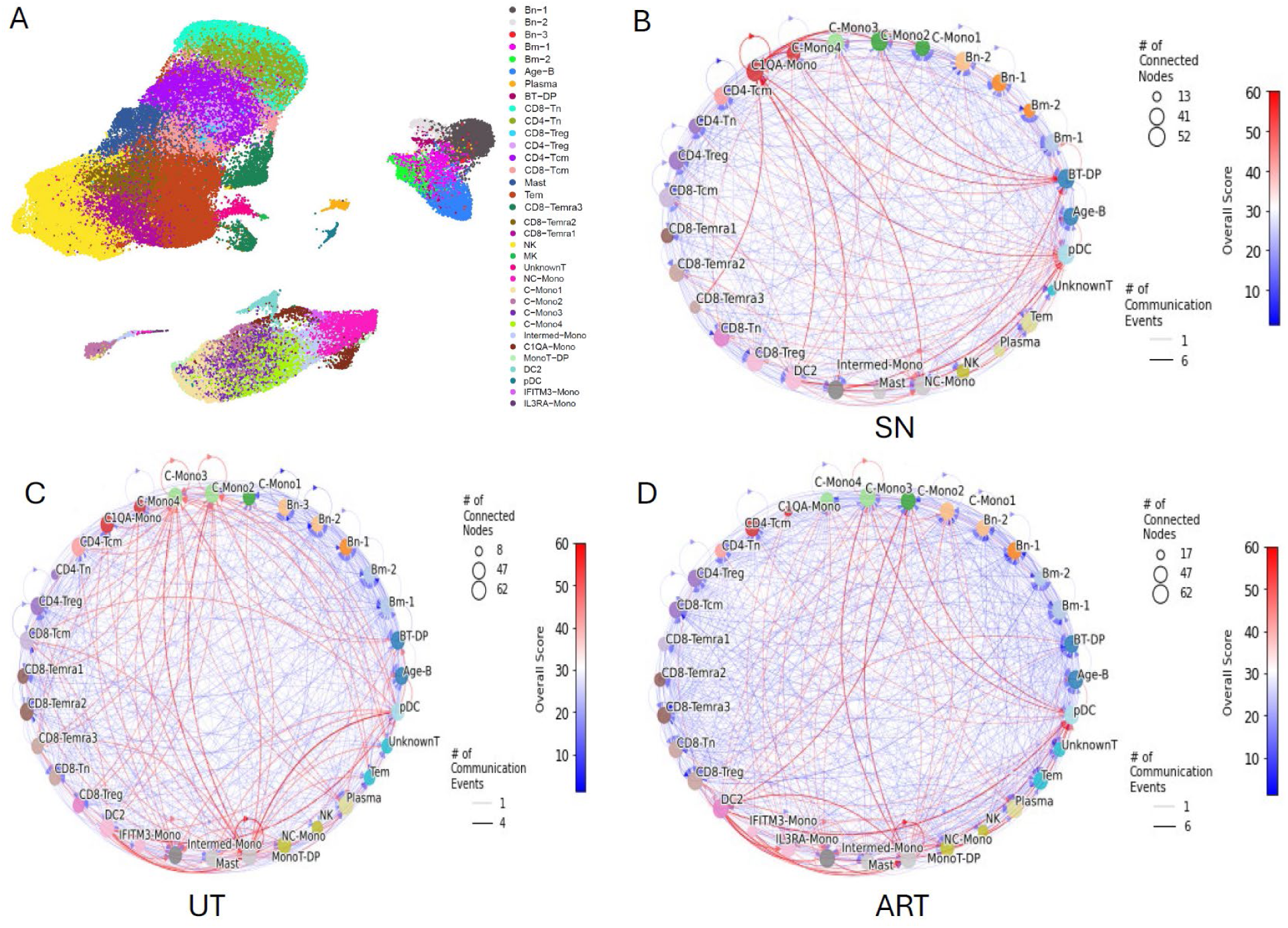
Identification of metabolite-based cell-cell communications of immune cell from PBMC on SN, UT and ART. A. UMAP plot of 34 sub-T, NK, B, monocytes and DC cell types of PBMC. B-D. These network plots illustrate the predicted cell-cell communication events based on metabolite interactions in PBMCs from SN(B), UT(C) and ART(D) using the MEBOCOST algorithm. Each node represents a distinct cell type, and the size of the node indicates the number of connected nodes. The edges between nodes represent communication events, where red lines denote higher levels of communication, and blue lines represent lower levels. The thickness of the lines reflects the number of communication events, with thicker lines indicating more frequent communications. The color gradient on the edges (red to blue) reflects the overall communication score which was calculated by the sum of -log10(FDR) of all metabolite-sensor communications between the sender and receiver types, with red indicating a higher score and blue indicating a lower one.

This analysis is conducted using MEBOCOST on scRNA-Seq data from PBMCs of SN, UT and ART patients, reveals key insights into the metabolite-based cell-cell communication landscape within the immune system. The communication network highlights extensive metabolic crosstalk among various immune cell types, providing a comprehensive overview of the interactions that underlie immune coordination.

The communication network for SN reveals that monocyte subtypes (e.g., C-Mono4, C1QA-Mono, Intermed-Mono) serve as central hubs of communication (Figure 5B, S5A). Monocytes, particularly C-Mono4 and C1QA-Mono, exhibit strong metabolic interactions with other immune cells, especially CD4+ and CD8+ T cells (Tcm, Tem, and Treg) as well as B cells (Bn-1, Bn-2). These connections suggest that monocytes may play a crucial role in regulating immune responses through metabolite exchange. In particular, C1QA-Mono, a subtype known for its involvement in phagocytosis and immune regulation, shows elevated communication scores with CD4+ T cells and B cells, highlighting its potential role in modulating adaptive immunity. CD4+ T cells, including both Tcm and Treg, are also actively engaged in metabolic communication, possibly influencing both effector and regulatory immune functions. The communication network shows significant interactions between dendritic cells (pDC, DC2) and NK cells, indicating potential collaboration in antigen presentation and innate immunity. Additionally, B cells (Bn-1 and Bn-2) display strong connections with multiple cell types, suggesting that metabolic support to B cells may be a critical factor in antibody production and memory B cell functions. Interestingly, NC-Mono, which are involved in patrolling vasculature and sensing environmental changes, show fewer connections compared to classical and intermediate monocytes. This might indicate a more specialized role for NC-Mono in metabolite-based communications, possibly linked to specific immune challenges.

In this analysis of PBMCs from UT HIV patients (Figure 5C, S5B), the predicted cell-cell communication network reveals a highly interconnected metabolic communication system, with monocytes, CD4+ and CD8+ T cells, and B cells serving as prominent hubs of interaction. The analysis shows that C-Mono and specialized monocytes like C1QA-Mono maintain significant metabolic interactions with both T cells and B cells, similar to the network observed in SN control.

A major difference between UT and SN control is the role of IFITM3-Mono, which shows higher communication activity in the UT network, particularly with monocyte and T cell subsets. This increased activity might reflect a heightened immune response, or an altered metabolic profile associated with the untreated HIV infection, suggesting that these monocytes are more metabolically active or involved in sensing viral signals. In contrast, NK cells and plasma cells display relatively less communication activity in the UT network compared to the SN network, potentially indicating a reduced innate immune response or altered antibody production mechanisms in untreated HIV patients. The NC-Mono, which are involved in patrolling and sensing environmental changes, also exhibit fewer interactions, suggesting a possible functional shift in this subtype in untreated HIV infection. Compared to the SN network, the B cells (Bn-1, Bn-2) in the UT network show a similar level of metabolic communication, implying that B cell functions remain relatively stable between seronegative and untreated conditions. However, dendritic cells (DC2, pDC) in the UT network exhibit more active communication with T cells, which could suggest an enhanced role in antigen presentation or immune regulation under untreated conditions.

The analysis of PBMCs from ART patients reveal a metabolic communication network that shows both similarities and distinctions from the networks observed in UT HIV patients and SN controls (Figure 5D, S5C). In ART patients, monocytes (particularly C-Mono, Intermed-Mono, and C1QA-Mono) continue to serve as key hubs of communication, maintaining strong metabolic interactions with T cells and B cells, similar to both the UT and SN networks. However, the overall communication intensity appears more balanced, with fewer highly concentrated red edges compared to the UT network, suggesting a partial normalization of immune function under ART. One notable difference in ART patients is the re-emergence of NK cells and plasma cells as active participants in the communication network, contrasting with their reduced role in untreated patients. NK cells exhibit more interactions with both T cells and monocytes in ART patients, indicating the restoration of some innate immune functions. This is particularly relevant as NK cells play a crucial role in controlling viral infections and mediating immune surveillance. IFITM3-Mono, which was highly active in the UT network, remains active in ART patients but shows slightly reduced communication intensity, suggesting that some of the heightened immune activation seen in untreated HIV is being dampened by ART. Similarly, NC-Mono, which had fewer interactions in UT patients, display more metabolic communications in ART patients, implying that ART may restore or preserve the patrolling functions of these monocytes. Compared to the SN network, the ART network still shows higher overall communication intensity, particularly among monocytes and T cells. However, the reduction in extreme communication events (red edges) compared to the UT network suggests that ART is helping to normalize immune metabolism, even though it does not completely restore the baseline communication seen in SN patients. The communication patterns in ART patients still reflect a somewhat dysregulated immune state, with ongoing high activity in specialized monocytes (C1QA-Mono, IFITM3-Mono) and B cells. When comparing all three conditions, the UT network exhibits the highest intensity of communication, particularly among monocytes and T cells, likely reflecting heightened immune activation due to uncontrolled HIV replication. In contrast, ART appears to mitigate this hyperactivation, restoring more balanced communication patterns similar to, but not fully equivalent to, the SN condition. ART patients show reduced but persistent metabolic crosstalk, indicating partial recovery of immune function, yet retaining signs of immune dysfunction or ongoing adaptation due to the chronic nature of HIV infection.

We further investigated the abundance of metabolites and the abundance of sensors across these patient groups (Figure 6). In the left-side panels of the figure, the top 10 metabolites show a distinct pattern in SN, UT, and ART patients. For SN patients, the metabolite profile reflects a baseline immune state, likely representing a balanced and homeostatic immune metabolism. These metabolites are associated with typical immune surveillance, cellular energy production, and basal signaling functions.

**Figure 6.**
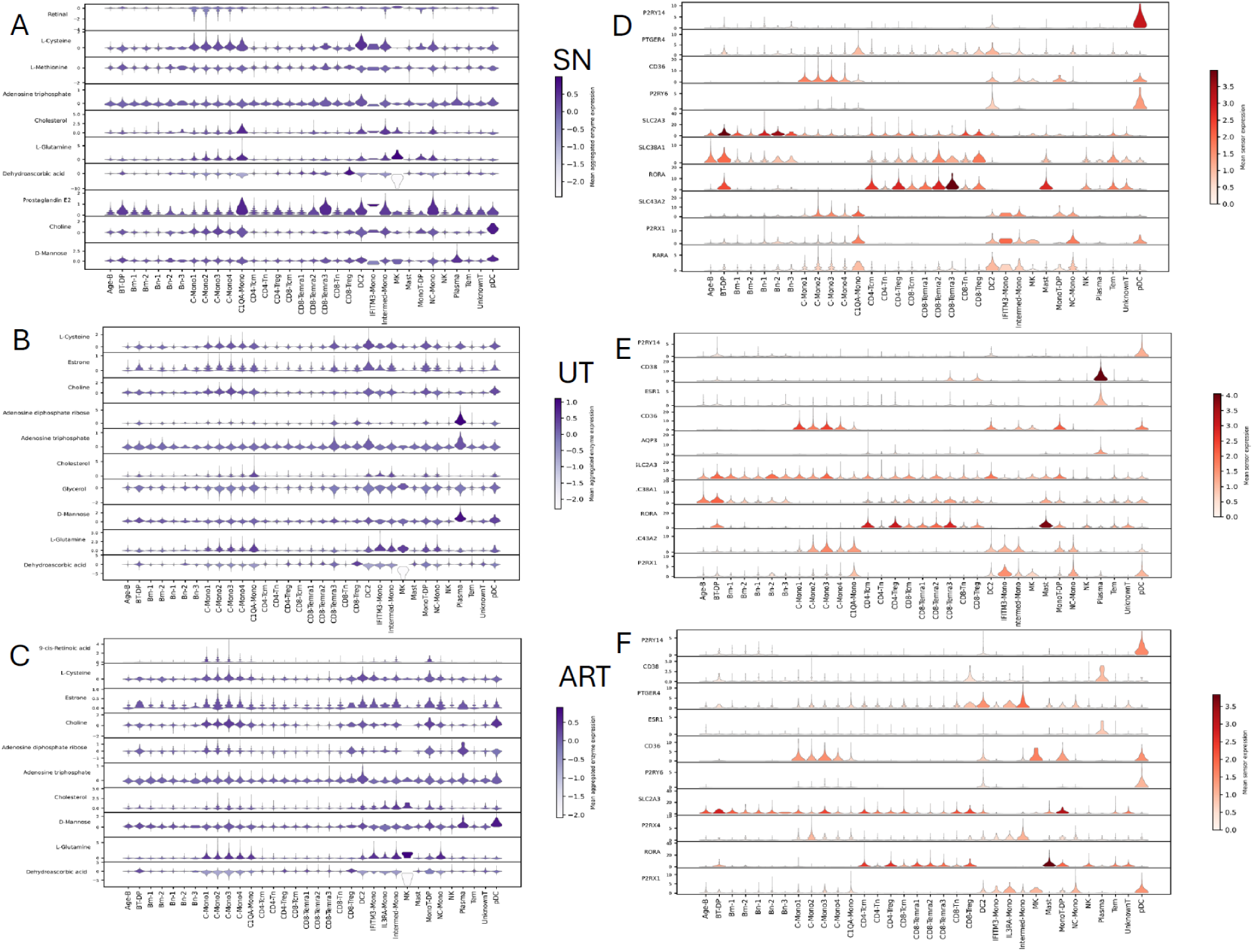
Violin plots of the abundance of top 10 representative metabolites (A, B and C) and sensors (D, E and F) across cell types in SN, UT and ART, respectively. Color bar shows Metabolite abundance(blue) and sensor abundance (red).

In contrast, UT patients show a drastically altered metabolite abundance profile. The top metabolites in this group are likely associated with heightened immune activation, viral sensing, and inflammatory responses. This increase in metabolite levels likely reflects the hyperactive immune environment in untreated HIV infection, where ongoing viral replication drives chronic immune stimulation and metabolic disruption. Metabolites involved in glycolysis, fatty acid metabolism, and oxidative stress may be elevated due to the heightened energy demands of immune cells fighting off infection.

ART patients show an intermediate metabolite profile compared to SN and UT groups. Although the levels of certain metabolites remain elevated relative to SN patients, the abundance is generally reduced compared to UT patients. This reflects partial metabolic normalization as a result of viral suppression under ART, although the immune cells may still retain some features of the hyperactivation seen in untreated infection.

The right-side panels of the figure highlight the top 10 sensor abundances. In SN patients, these sensors reflect a balanced state of immune monitoring, with typical sensors involved in detecting metabolic and immune signals. UT patients, however, show a shift towards sensors associated with immune activation and stress, which may include those involved in recognizing viral components or damage-associated signals. The elevated sensor abundance in UT patients is likely a result of chronic immune stimulation and the immune system’s attempt to control viral replication.

For ART patients, the sensor profile is somewhat restored towards a more normalized state, although it does not fully return to the levels observed in SN patients. Some sensors associated with immune activation and viral recognition remain elevated, reflecting the chronic nature of HIV infection, even when controlled by ART. This may suggest that while ART reduces viral load, the immune system remains in a heightened state of alertness, likely due to persistent viral reservoirs or low-level inflammation.

In conclusion, the comparison between SN, UT, and ART samples demonstrates that HIV infection significantly alters both metabolite and sensor abundance in immune cells. UT patients exhibit the most dramatic shifts, reflecting immune hyperactivation and metabolic dysregulation. ART partially normalizes these profiles but does not fully restore them to the baseline observed in SN patients. This ongoing dysregulation in ART patients may contribute to immune exhaustion or persistent inflammation, factors that are important in the long-term management of HIV despite effective viral suppression.

The metabolite and sensor changes among SN, UT, and ART patients reveal important insights into the immune-metabolic landscape affected by HIV infection and its treatment. In SN patients, the top metabolites reflect a balanced, homeostatic immune metabolism. These metabolites are likely involved in normal cellular respiration, lipid metabolism, and immune signaling, maintaining baseline immune functions such as surveillance and regulation. In contrast, UT patients exhibit significant changes in metabolite abundance. Metabolites related to glycolysis, fatty acid oxidation, and inflammatory pathways are often upregulated in UT patients, indicative of chronic immune activation and metabolic stress. This metabolic shift reflects the high energy demands of immune cells continuously battling HIV replication, alongside elevated production of ROS and inflammatory mediators.

In ART patients, while many metabolite levels are reduced compared to the UT group, they do not fully normalize to the levels seen in SN individuals. Some of the elevated metabolites in ART patients suggest lingering immune activation or metabolic remodeling due to chronic infection, despite viral suppression. This suggests that ART mitigates the hypermetabolic state but doesn’t completely restore metabolic homeostasis.

In SN individuals, sensor abundance corresponds to normal immune surveillance, with key immune sensors functioning to detect regular metabolic and pathogen-associated signals. In UT patients, sensors linked to immune stress, viral sensing (such as pattern recognition receptors), and inflammation is highly upregulated. This mirrors the heightened immune response against HIV, as cells ramp up their detection capabilities in response to viral infection.

Under ART, sensor abundance decreases compared to UT patients, reflecting partial immune recovery. However, certain sensors remain elevated, pointing to residual immune alertness and an inability to fully return to a baseline state. This continued dysregulation could be associated with immune exhaustion or low-grade chronic inflammation, hallmarks of treated but persistent HIV infection. Thus, while ART significantly curtails viral replication, it does not entirely reverse the immune system’s heightened state of vigilance or metabolic disruption.

## 4. Discussion

Understanding the metabolic changes in immune cells during infections, particularly HIV, is crucial for comprehending immune responses. Metabolic reprogramming is essential for effective immunity, but chronic infections can exploit or exhaust these pathways, leading to dysfunction. Insights into these metabolic shifts open avenues for therapeutic interventions, targeting metabolic pathways to enhance immune function or inhibit pathogen survival. However, few studies have been conducted on the metabolic dynamics of immune cells during HIV infection and after antiretroviral therapy using scRNA-Seq.

Cell-cell communication is a fundamental mechanism that coordinates cellular activities in development and disease. Besides protein ligands, metabolites are another major type of signaling molecules mediating cell-cell communications, extensively studied through experimental approaches. By leveraging the newly developed computational tool MEBOCOST, we can study metabolite-based intercellular communications using single-cell RNA-seq data.

In this study, we separated PBMCs into T and NK cells, monocytes and DCs, and B cells. We then clustered these cell populations into subtypes. For each subtype, we investigated all metabolic pathways by comparing UT and ART patients with seronegative controls. Metabolic pathways, including energy production and utilization, lipid metabolism, amino acid metabolism, glycosylation and glycan biosynthesis, and oxidative stress and antioxidant defense, are reprogrammed in different immune cells.

Next, we examined metabolite-mediated cell-cell communications in SN, UT, and ART groups. In untreated HIV patients, the predicted cell-cell communication network reveals a highly interconnected metabolic communication system, with monocytes, CD4+ and CD8+ T cells, and B cells serving as prominent hubs of interaction. The analysis shows that C-Mono and specialized monocytes like C1QA-Mono maintain significant metabolic interactions with both T cells and B cells, similar to the network observed in seronegative patients. However, the communication patterns in UT patients are more extensive, with denser interactions and more frequent communication events, represented by thicker and more abundant red edges.

In ART patients, the metabolic communication network shows both similarities and distinctions from the networks observed in untreated HIV patients and seronegative controls. The overall communication intensity appears more balanced, with fewer highly concentrated red edges compared to the UT network, suggesting a partial normalization of immune function under ART. While ART significantly impacts the immune-metabolic landscape, reducing the intense communications seen in untreated patients, the immune system in ART patients remains more active than in SN controls, indicating that HIV infection continues to alter immune metabolism even under treatment.

This study provides valuable insights into the metabolic changes and metabolite-mediated cell communication in immune cells across various immune diseases. These findings highlight the complex interplay between metabolism and immune function and underscore the potential for metabolic interventions to improve treatment outcomes. By targeting specific metabolic pathways, it is possible to develop novel therapeutic strategies that enhance immune function, control disease progression, and improve the quality of life for individuals living with immune diseases.

## Abbreviations

UT: untreated HIV-1 individuals
ART: HIV-1-infected individuals receiving antiretroviral therapy (ART)
SN: seronegative controls
scRNA-Seq: single cell RNA sequencing
NK: natural killer cell
MK: megakaryocyte
pDC: plasma dendritic cell.
Tn: naïve T cell
Tcm: central memory T cell
Treg: regulatory T cell
Tem: effector memory T cells
Temra: terminally differentiated effector memory T cells
C-Mono: classical monocytes
NC-Mono: non-classical monocytes
Intermed-mono: intermediate monocytes
Bn: naïve B cell
Bm: memory B cell
BT-DP: B and T double positive cell.

## Author Contributions

D.W. and P.X. conceived the project. P. X. curated the scRNA-seq data and performed the computational analysis. D.W. and P. X. analyzed and interpreted the results. P. X. drafted and edited the manuscript. All authors read and edited the manuscript.

---

**Figure S1.**
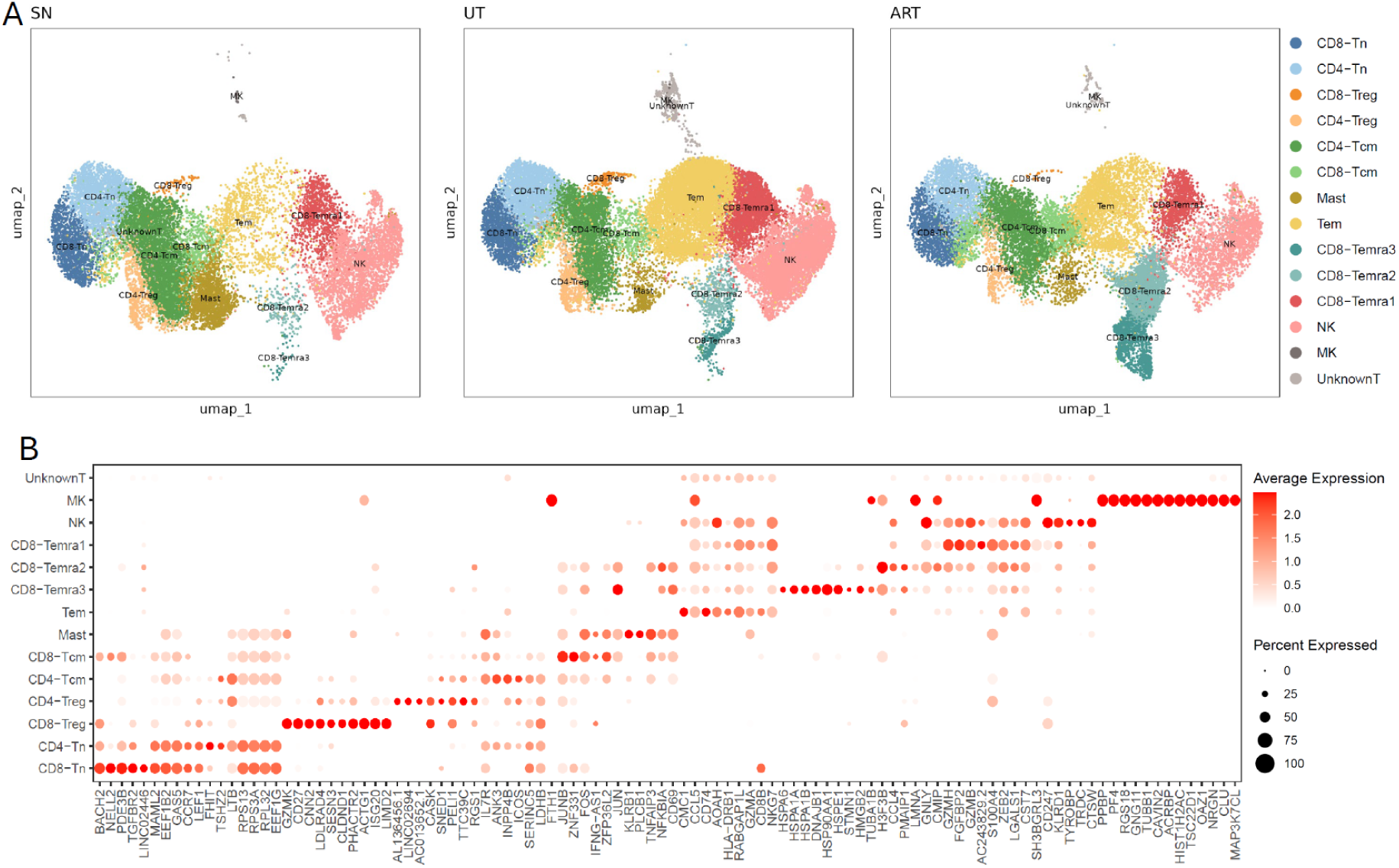
(A) UMAP plots of 8 T and NK cell subpopulations of PBMC from SN, UT and ART patients. (B) Dotplot showing the top ten markers predicted for each sub cell type.

**Figure S2.**
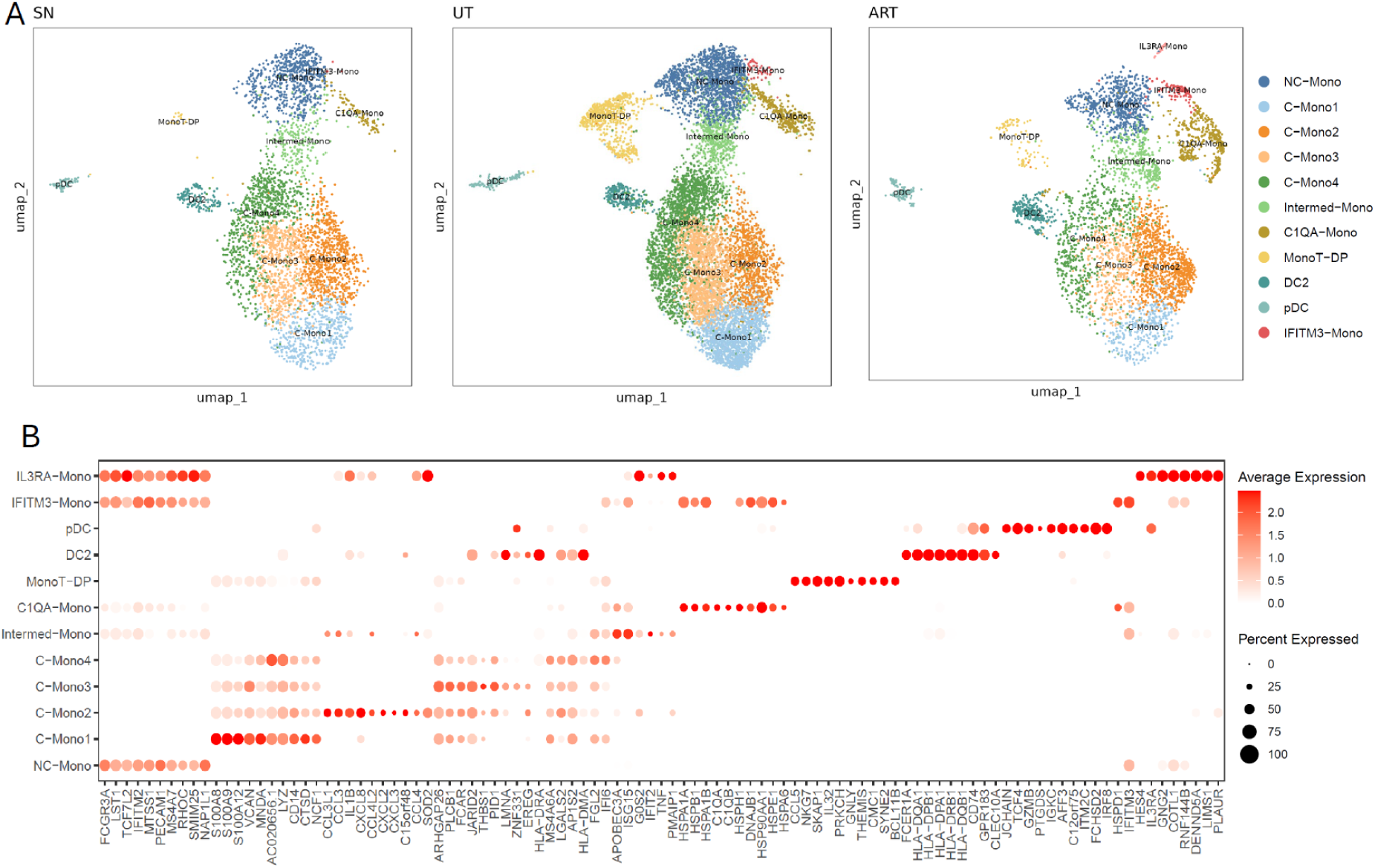
(A) UMAP plots of 11 Monocytes and DC cell subpopulations of PBMC from SN, UT and ART patients. (B) Dotplot showing the top ten markers predicted for each sub monocytes and DC cell type.

**Figure S3.**
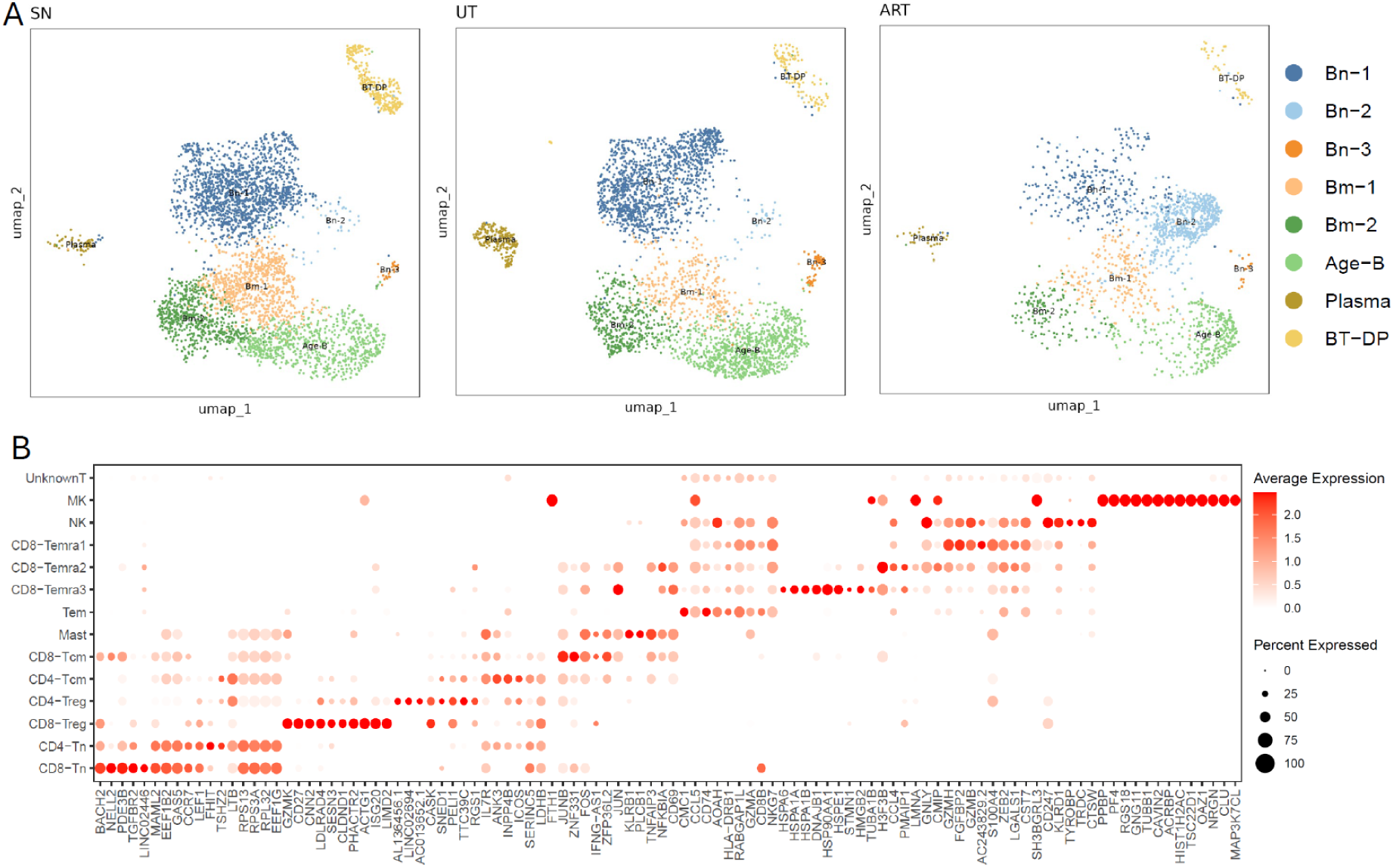
(A) UMAP plots of 8 B cell subpopulations of PBMC from SN, UT and ART patients. (B) Dotplot showing the top ten markers predicted for each sub-B cell type.

**Figure S4.**
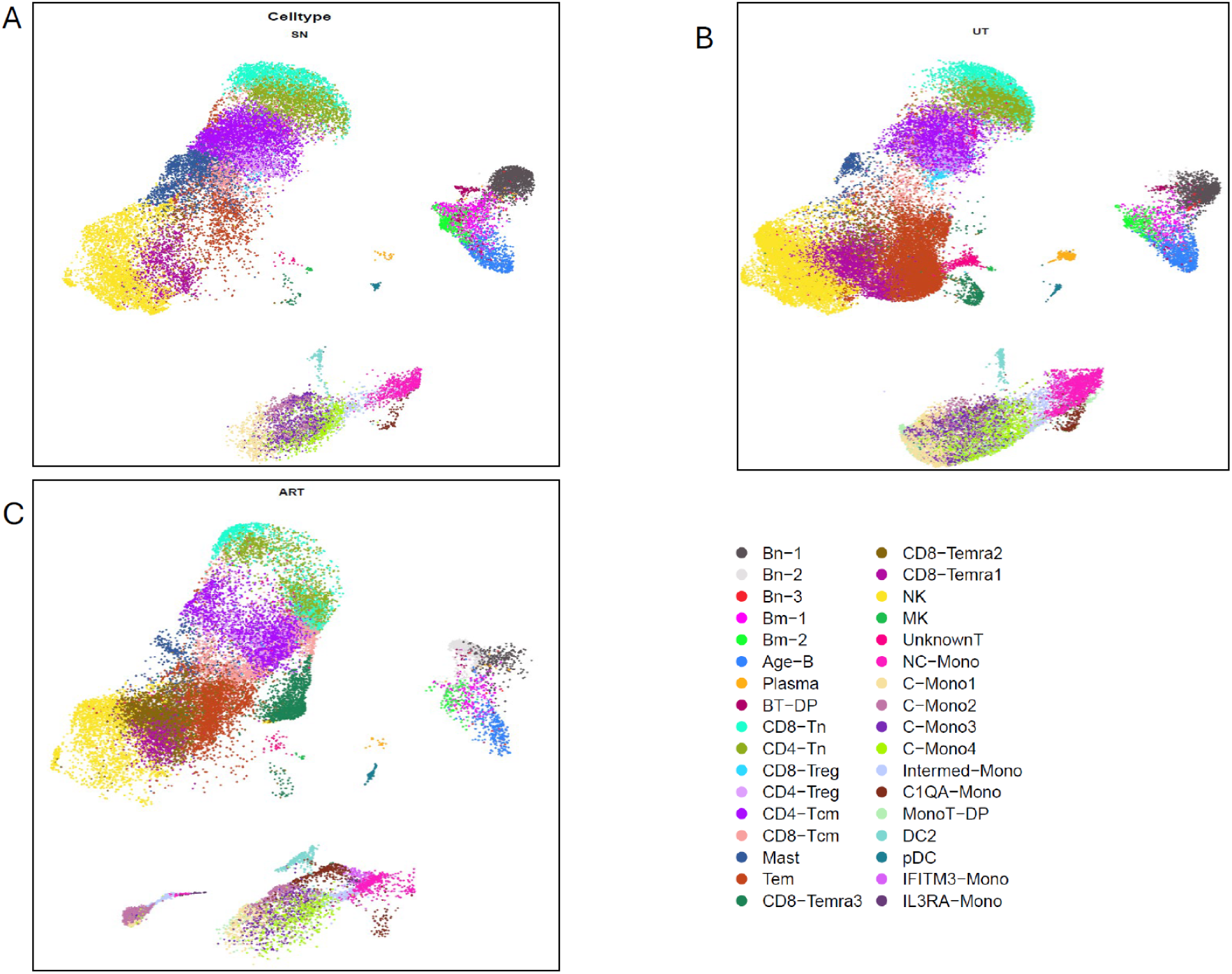
UMAP plot of 34 sub-T, NK, B, monocytes and DC cell types of PBMC on SN(A), UT(B) and ART(C), respectively.

**Figure S5.**
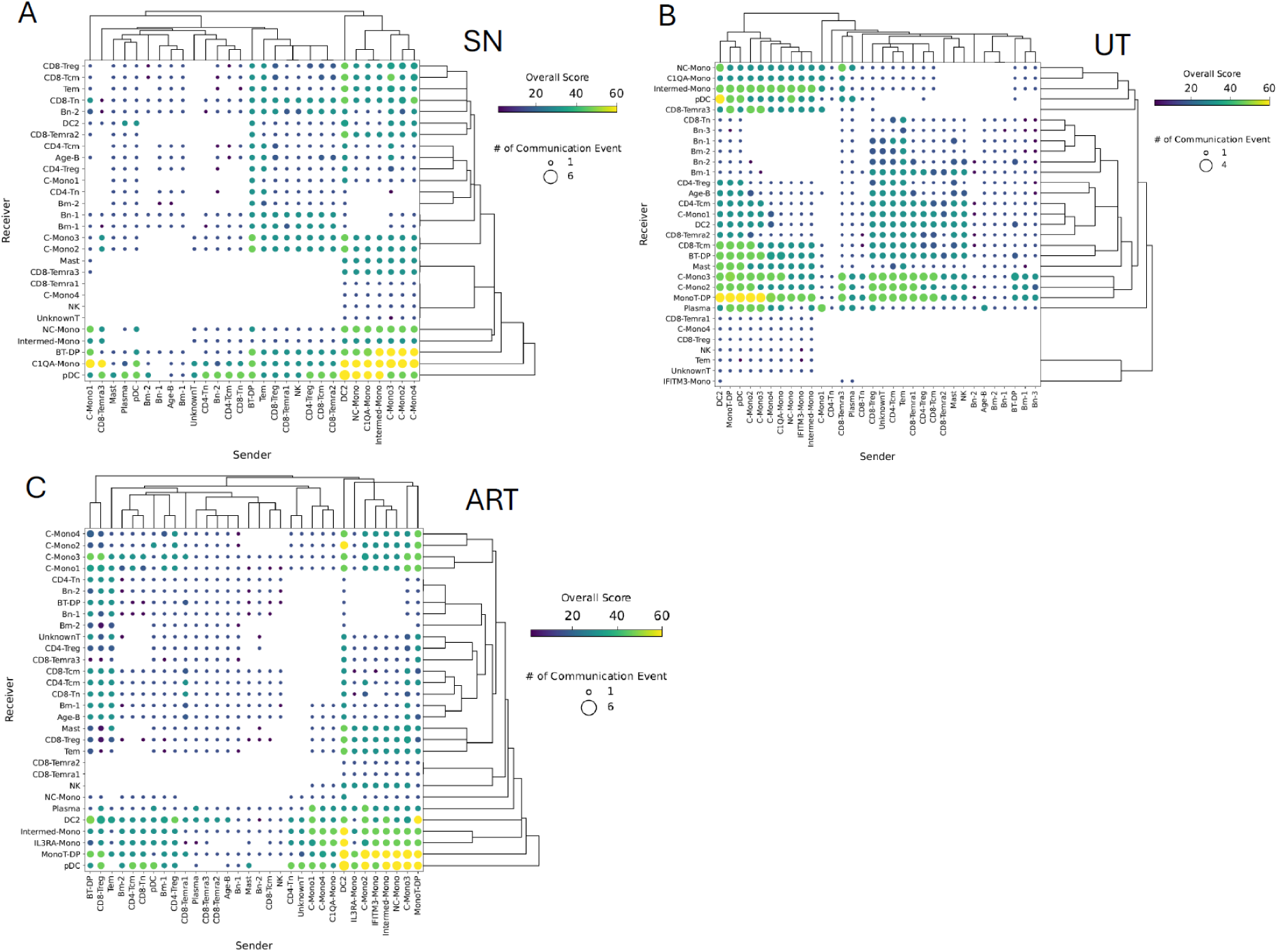
Dot plots show the communication of sub cell types between sender and receiver in SN(A), UT(B) and ART(C) patients.

